# Phenotypic analysis of 11,125 trio exomes in neurodevelopmental disorders

**DOI:** 10.1101/2025.03.11.642649

**Authors:** Shiva Ganesan, Sarah M. Ruggiero, Shridhar Parthasarathy, Peter D. Galer, David Lewis-Smith, Ian McSalley, Stacey R. Cohen, Laina Lusk, Anna J. Prentice, Jillian L. McKee, Manuela Pendziwiat, Lacey Smith, Yvonne Weber, Heather C. Mefford, Annapurna Poduri, Ingo Helbig

## Abstract

Genomic sequencing is widely used to identify causative genetic changes in neurodevelopmental disorders, such as autism, intellectual disability, and epilepsy. Most neurodevelopmental disorders also present with diverse clinical features, and delineating the interaction between causative genetic changes and phenotypic features is a key prerequisite for developing personalized therapies. However, assessing clinical features at a scale that parallels genomic sequencing remains challenging. Here, we standardize phenotypic information across 11,125 patient-parent trios with exome sequencing data using biomedical ontologies, analyzing 674,767 phenotypic terms. We find that individuals with *de novo* variants in 69 out of 261 neurodevelopmental genes exhibit statistically significant clinical similarities with distinct phenotypic fingerprints. We also observe that phenotypic relatedness follows a gradient, spanning from highly similar to dissimilar phenotypes, with intra-gene similarities suggesting clinically distinct subgroups for seven neurodevelopmental genes. For most genetic etiologies, only a small subset of highly phenotypically similar individuals carried *de novo* variants in the same gene, highlighting the heterogeneous and complex clinical landscape of neurodevelopmental disorders. Our study provides a large-scale overview of the dynamic relationship between genotypes and phenotypes in neurodevelopmental disorders, underscoring how the inherent complexity of these conditions can be deciphered through approaches that integrate genomic and phenotypic data.

## Introduction

Neurodevelopmental disorders affect 1-2% of children worldwide and result from a diverse range of etiologies including brain injuries, structural brain malformations, and genetic variants^1^. Over the last two decades, the identification of causative genetic etiologies has led to a paradigm shift in understanding these conditions. Up to 20% of individuals with autism, intellectual disability, and epilepsy have causative genetic changes identified through novel genomic techniques such as massively parallel sequencing^2^. Given the overall frequency of monogenic causes among neurodevelopmental disorders, disease-modifying therapies are becoming increasingly available, including antisense oligonucleotide (ASO) therapies and adeno-associated virus (AAV) based gene therapies^3,4^.

The development of precision medicine approaches and assessment of their efficacy critically depend on a detailed understanding of the clinical features associated with specific genetic causes^5^. Historically, small case series have provided critical insights into the clinical spectrum and associated features of genetic neurodevelopmental disorders^6,7^. However, with over 250 genetic causes of neurodevelopmental disorders identified, the capacity to perform traditional clinical studies has emerged as a major bottleneck^8^. Given the growing number of neurodevelopmental genes, it has become impossible to generate sufficient phenotypic detail to characterize each disease entity. This presents challenges both with designing outcome measures for clinical trials and providing accurate medical guidance for individuals affected by genetic neurodevelopmental disorders^5^.

Novel methods for large-scale phenotype analysis offer alternative approaches for understanding the clinical spectrum within a wide range of rare disorders, including neurodevelopmental disorders^9–13^. Previous studies have introduced novel approaches through analyzing phenotypic data from large patient cohorts using clinical data harmonized through biomedical dictionaries^9,10^. These approaches have been sufficiently powered to explore the range of clinical features within a broad range of conditions and highlight previously unrecognized disease patterns within broader patient populations^12,13^.

Large-scale phenotyping studies combined with genomic data have historically been constrained by the heterogeneous documentation of clinical features. This roadblock has been addressed by emerging tools for standardizing phenotypic terminology, including biomedical dictionaries and ontologies^14^. One of the most commonly used dictionaries in clinical genomic testing and research is the Human Phenotype Ontology (HPO), a hierarchical dictionary of more than 13,000 clinical terms designed to facilitate computational approaches^15,16^. This roadblock has been addressed by emerging tools for standardizing phenotypic terminology, including biomedical dictionaries and ontologies^14^. One of the most commonly used dictionaries in clinical genomic testing and research is the Human Phenotype Ontology (HPO), a hierarchical dictionary of more than 13,000 clinical terms designed to facilitate computational approaches^15,16,10^. HPO annotations combined with statistical methods have previously been used to discover novel genetic etiologies in neurodevelopmental disorders^6^, map the phenotypic landscape of known genetic etiologies^11^, and address the longitudinal trajectories within a subset of genetic neurodevelopmental disorders^9,13^. In addition, biomedical dictionaries such as the HPO lend themselves to machine learning approaches that have successfully predicted genetic etiologies based on early clinical features^9,12^.

Here, we analyze 674,767 clinical phenotypic annotations in 11,125 individuals with genetic neurodevelopmental disorders, integrating exome sequencing (ES) data with phenotypic annotations. We find that this approach allows us to delineate the complex interactions between genetic etiologies and phenotypic features, including measures of clinical resemblance across disorders, disease subgroups, and the pattern of causative genetic alterations in groups of individuals that are highly clinically similar.

## Results

### Harmonization of multiple cohorts using an HPO-based framework

To systematically review the interplay between genetic and clinical features, we aggregated clinical and genomic data from four different cohorts: a cohort from a pediatric tertiary care center obtained through diagnostic sequencing (n=192) with a median of 12 clinical features annotated per individual, the Epilepsy Phenome-Genome Project (EPGP; n=335) with a median of nine clinical features^17^, the Deciphering Developmental Disorders study (DDD; n= 13,424) with a median of six clinical features^18^, the EuroEPINOMICS-RES cohort (n=319) with a median of 10 clinical features^19^, and the Children’s Hospital of Philadelphia (CHOP) Birth Defects Biorepository (n=623) with a median of eight HPO terms. Using the framework of the HPO^15^, we harmonized clinical data across all four cohorts. After data integration, we retrieved a total of 112,733 clinical annotations in 14,893 individuals, including 11,125 individuals with available trio ES data. The clinical annotations consisted of 4,686 unique HPO terminologies, including global developmental delay (HP:0001263, f=26.1%), delayed speech and language development (HP:0000750, f=15.6%), and seizures (HP:0001250, f=11.9%) as the most common. Among these, 1,952 out of 4,686 annotations (41.7%) were present in two or less individuals, highlighting a sparsity of clinical annotations that has also been observed in prior studies, as well as the presence of clinical features unique to particular genetic neurodevelopmental disorders^13^. The amount of phenotypic information per individual was also variable, with the number of annotations per individual ranging from 1 to 64 terms, with a median of six clinical terms per individual. Leveraging the hierarchal structure of HPO to infer higher-level terms, we expanded the dataset to derive a total of 674,767 clinical annotations, which included 5,707 distinct terms with a median of 41 phenotypic features per individual (range of 3-251 clinical features; **Fig. 1**). The inclusion of more general phenotypic features is referred to as propagation and has previously been shown to aid the harmonization of clinical information by leveraging the tree-like structure of the underlying biomedical ontology^9^.

**Fig. 1:**
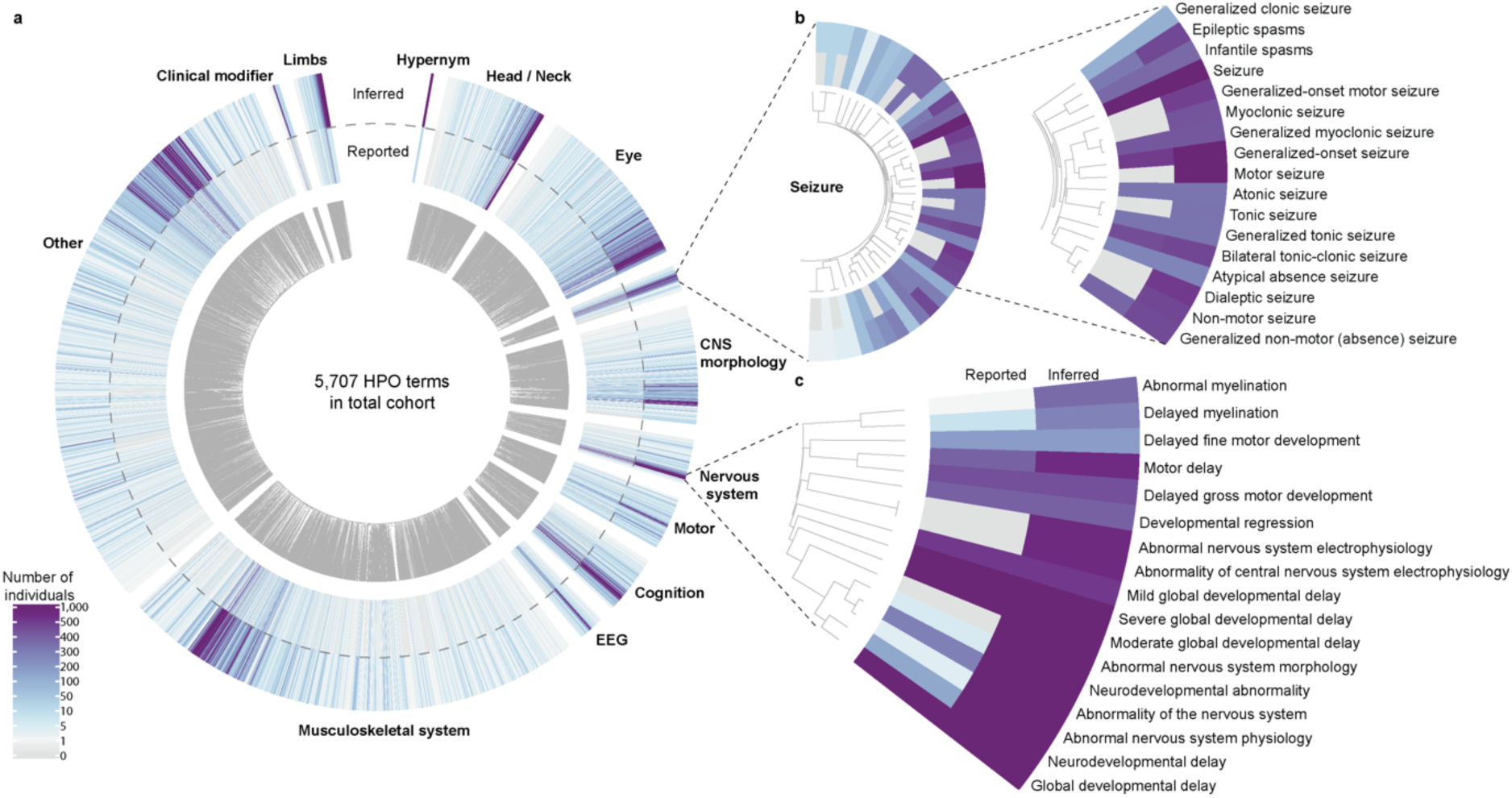
Harmonization of clinical features using HPO. **a**, Clinical features for 14,893 individuals mapped to 4,686 distinct HPO terms, yielding 5,707 inferred HPO clinical features after harmonization. **b**, Seizure subontology with 55 reported and 81 inferred terms across 3,324 individuals, with the most frequent clinical annotation terms highlighted. **c**, The Abnormality of nervous system HPO branch, highlighting the importance of harmonization. The process of harmonization using HPO lead to nearly six-fold increase in the total data points, increasing the median number of terms per individual from 6 to 41.

### Genomic analysis identified 261 distinct genetic etiologies with *de novo* variants

After harmonizing phenotypic data across all four cohorts, we sought to identify potentially disease-causing variants among 11,125 available trio exomes, including proband and parental data. We utilized a standardized pipeline across all samples for variant calling^20^, annotation, and assessment of *de novo* variants^21^, the most common genetic mechanism observed in neurodevelopmental disorders^22,23^. We applied standard filters to the analysis of genomic data, including heterozygous variants, allele read depth > 10, allele balance between 0.25 and 0.75, and exclusion of population variants present in gnomAD v4.1^22,23^. Applying these criteria to the existing trio ES data, we identified a total of 1,969 *de novo* variants, including 261 distinct genetic etiologies with *de novo* variants in two or more individuals (**Extended Data Table 1**).

The most frequent genetic etiologies observed in the combined cohort included *KMT2A* (n=22), *KCNQ2* (n=20), *STXBP1* (n=17), *MECP2* (n=16), and *PPP2R5D* (n=16). A total of 18 genetic etiologies had *de novo* variants detected in >10 individuals, and 130 genetic etiologies were identified in only two individuals. We also identified a total of 81 recurrent *de novo* variants in 199 individuals among 64 distinct genetic etiologies present in two or more individuals (**Extended Data Table 2**). The most frequent recurrent *de novo* variants included *PPP2R5D*:p.Glu198Lys (n=12), *ADNP*:p.Tyr719Ter (n=6), *HNRNPH2*:p.Arg206Trp (n=6), *MECP2*:p.Arg145Cys (n=5), and *AP2M1*:p.Arg710Trp (n=4). Using denovolyzeR^24^, we identified 155 out of 261 genetic etiologies significantly enriched for *de novo* variants within the joint cohort, providing statistical evidence supporting the involvement of these 155 genetic etiologies in neurodevelopmental disorders. The framework to identify disease-causing variants by comparing observed versus expected frequency of *de novo* variants is well established^23^. We used this framework to assess the significance for each genetic etiology based on genomic features only, referred to here as the genomic significance.

### Analysis of phenotypic features enables assessment of phenotypic similarity

We next evaluated phenotypic similarities for each genetic etiology with the aim to contrast significance based on the frequency of *de novo* variants (genomic significance) with significance based on clinical resemblance for a given genetic etiology (phenotypic significance). We performed a phenotypic similarity analysis, an established method to assess clinical relatedness, which utilizes the structure of the HPO to compare the intersection of phenotypic features between individuals^6,12,15,25^. In brief, a phenotypic similarity analysis of a subset of individuals assesses whether clinical features observed in those individuals are more related than expected by chance. We computed similarity score based on the most informative common ancestor concept, assessing pairwise similarity of individuals using the combinatorial comparison of specific phenotypes shared between individuals. Methods to perform phenotypic similarity analyses based on HPO terms have previously been developed and can be performed using various algorithms^26^ (**Extended data Fig. 1)**.

We identified 69/261 genes with nominally significant phenotypic similarity, representing genes where phenotypes in individuals with *de novo* variants were more phenotypically related than expected by chance (**Fig. 2a)**. *SCN1A* (n=14), *AP2M1* (n=4), and *DNM1* (n=6) were the most phenotypically similar genes. For *SCN1A*, prominent features driving phenotypic similarity were bilateral tonic-clonic seizures (HP:0002069; freq = 80%; p-val= 8.8x10^-16^), focal clonic seizures (HP:0002266; freq = 53%; p-val= 2.7x10^-15^), and febrile seizures (HP:0002373; freq = 67%; p-val= 1.9x10^-14^). In contrast, for *AP2M1*, the most prominent features were interictal epileptiform activity (HP:0011182; freq = 100%; p-val= 7.35x10^-6^), atonic seizures (HP:0010819; freq = 75%; p-val= 1.4x10^-5^), and EEG with spike-wave complexes (HP:0010850; freq = 75%; p-val= 2.4x10^-5^). For *DNM1*, myoclonic seizures (HP:0032794; freq = 83%; p-val= 1.06x10^-7^), EEG with spike-wave complexes < 2.5 Hz (HP:0010847; freq = 66%; p-val= 1.5x10^-7^), and atypical absence seizures (HP:0007270; freq = 66%; p-val= 1.89x10^-7^) were the driving features.

**Fig. 2:**
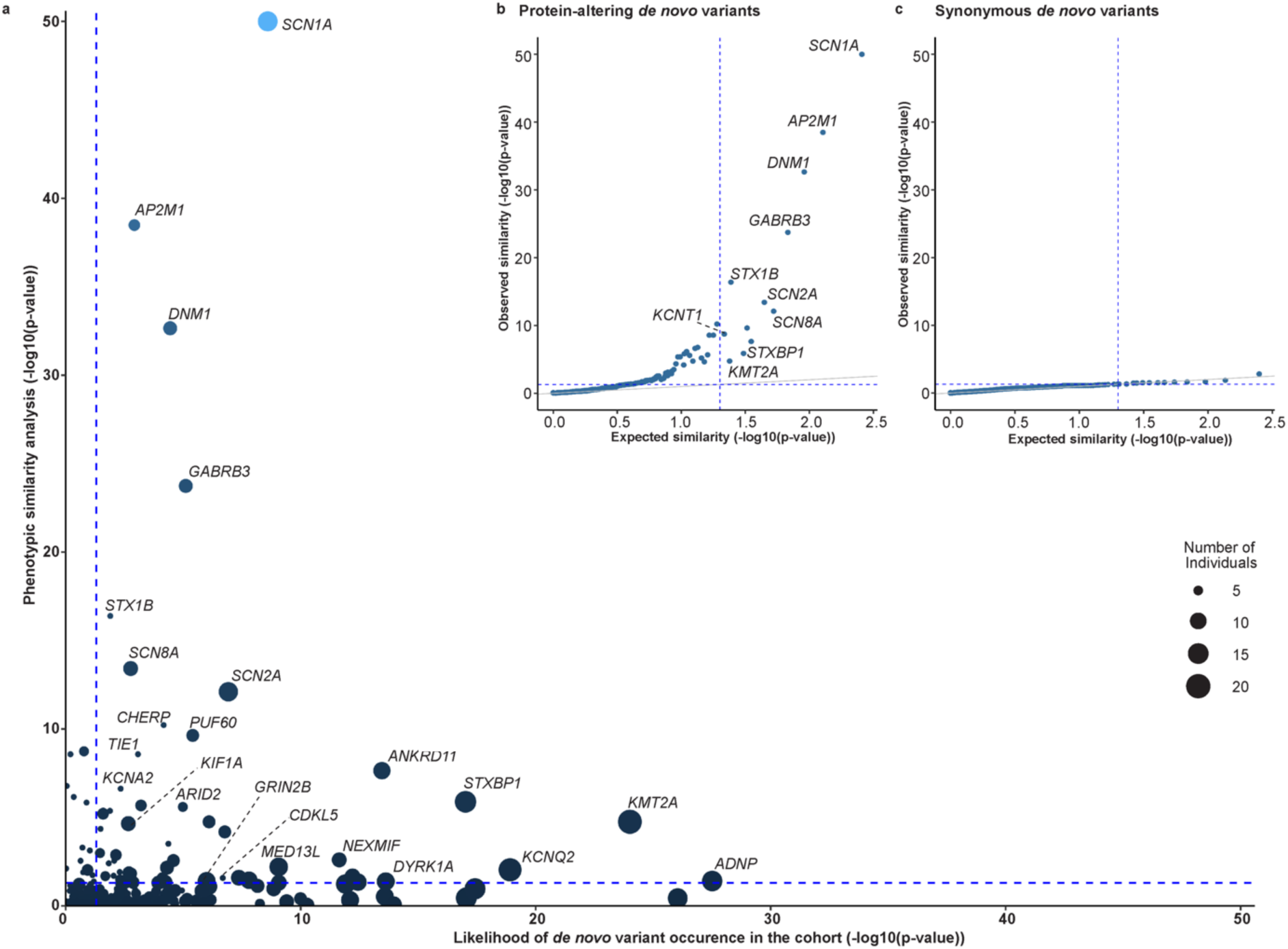
Comparison of statistical significance of *de novo* variant occurrence and phenotypic similarity. **a**, Comparison of statistical significance of 261 genes with *de novo* variants, integrating both genetic and phenotypic evidence. The x-axis represents the likelihood of the occurrence of *de novo* variants in the cohort computed using denovolyzeR, while y-axis represents the phenotypic similarity significance calculated through permutation analysis. We identified 69 genes significant based on phenotypic similarity, among which 15 genetic etiologies, including *SCN2A* and *KCNA2*, where phenotypic evidence exceeded genetic evidence. **b**, qqplot comparing the expected and observed phenotypic similarity for genetic etiologies with protein-altering *de novo* variants. Several genes, including *SCN1A*, *AP2M1*, and *DNM1* show higher phenotypic significance than expected. **c**, To assess algorithm performance, we computed phenotypic similarity for genes with synonymous *de novo* variants. The observed similarity remained low, validating the robustness of our method and are unlikely due to random chance and serve as a negative control.

### Genetic etiologies demonstrate variable patterns of phenotypic and genomic significance

We next compared phenotypic significance to genomic significance in all 261 genetic etiologies examined (**Fig. 2a)**, comparing the -log_10_(p-value) for both analyses. We found that a subset of 14 genes, including *MECP2*, *SYNGAP1,* and *KCNB1*, had an excess of *de novo* variants but low phenotypic similarity. Accordingly, these genetic etiologies, while relatively common, show highly diverse phenotypes that cannot be clearly distinguished from the larger cohort. Conversely, a subset of 37 genes had a more substantial phenotypic significance than genomic significance, highlighting genetic etiologies in the cohort that had strong phenotypic resemblance, even if they were rarely observed. These genes included *SCN1A*, *AP2M1*, and *DNM1*. We had previously predicted the existence of genetic etiologies identifiable through strong clinical resemblance rather than a burden of *de novo* variants^13^. The current analysis emphasizes that these genetic etiologies are hidden in large, heterogeneous datasets and can be identified through phenotypic similarity approaches.

No single genetic etiology stood out with simultaneous strong genomic and phenotypic significance; an observation that seems counterintuitive. It is possible that the datasets examined in our study are depleted for genetic disorders that are both common and recognizable, such as tuberous sclerosis complex and Angelman syndrome, as these disorders may be either subject to population-based screening due to their relative frequency or diagnosed via targeted testing due to their unique clinical manifestations^27–29^.

### Phenotypic similarity algorithms converge in large datasets

Finally, we examined the impact of various similarity algorithms with respect to their ability to identify genetic etiologies with clinical similarity. We had previously reasoned that various algorithms may differentially emphasize selected features of the HPO tree, such as density of sub-branches. However, in our current cohort, we observe that three different algorithms largely generate comparable results, when assessing the significance and ranking of specific genes (**Extended data Fig. 1)**. Accordingly, prior observations suggesting a divergence of results of phenotypic similarity algorithms may have been falsely exaggerated by limited sample sizes. Compared to individuals with synonymous variants, *de novo* protein-disrupting variants displayed a pattern of higher-than-expected similarity across all algorithms, with several genes consistently displaying the highest phenotypic similarity across multiple algorithms (**Fig. 2b-2c**).

### Analysis of phenotypes in recurrent variants reveals “diseases within diseases”

While clinical homogeneity can be detected at the gene level, the same method can be extended to recurrent *de novo* variants. Among the 81 recurrent *de novo* variants detected in our cohort, 26 showed significant phenotypic similarity (**Fig. 3**). For three genes (*KIF1A*, *SLC6A1*, *SRCAP*), phenotypic similarity of recurrent variants exceeded gene-level median similarity by two-fold, and for five genes (*SCN2A*, *SMARCA2*, *KCNQ2*, *ADNP*, *TCF7L2*), phenotypic similarity of recurrent variants surpassed gene-level similarity by three-fold. The most prominent phenotypic resemblance was seen for *SCN2A:*p.Arg853Gln, with EEG with focal sharp waves (HP:0011196), infantile spasms (HP:0012469), and developmental regression (HP:0002376) representing the most prominent phenotypes driving clinical relatedness. For *SLC6A1*, the p.Phe465Ser variant showed a two-fold more prominent phenotypic similarity driven by seizures (HP:0001250), bilateral tonic-clonic seizures with focal onset (HP:0007334), and growth delay (HP:0001510). For *KCNQ2:*p.Gly281Arg, we observed a three-fold increase in phenotypic similarity resulting from shared clinical features including encephalopathy (HP:0001298), generalized tonic seizures (HP:0010818), and EEG with generalized epileptiform discharges (HP:0011198). The most common recurrent variant in the cohort was *PPP2R5D:*p.Glu198Lys (n=12). Although the gene did not reach statistical significance, this recurrent variant exhibited higher similarity and phenotypic significance driven by macrocephaly (HP:0000256), and hypotonia (HP:0001252).

**Fig. 3:**
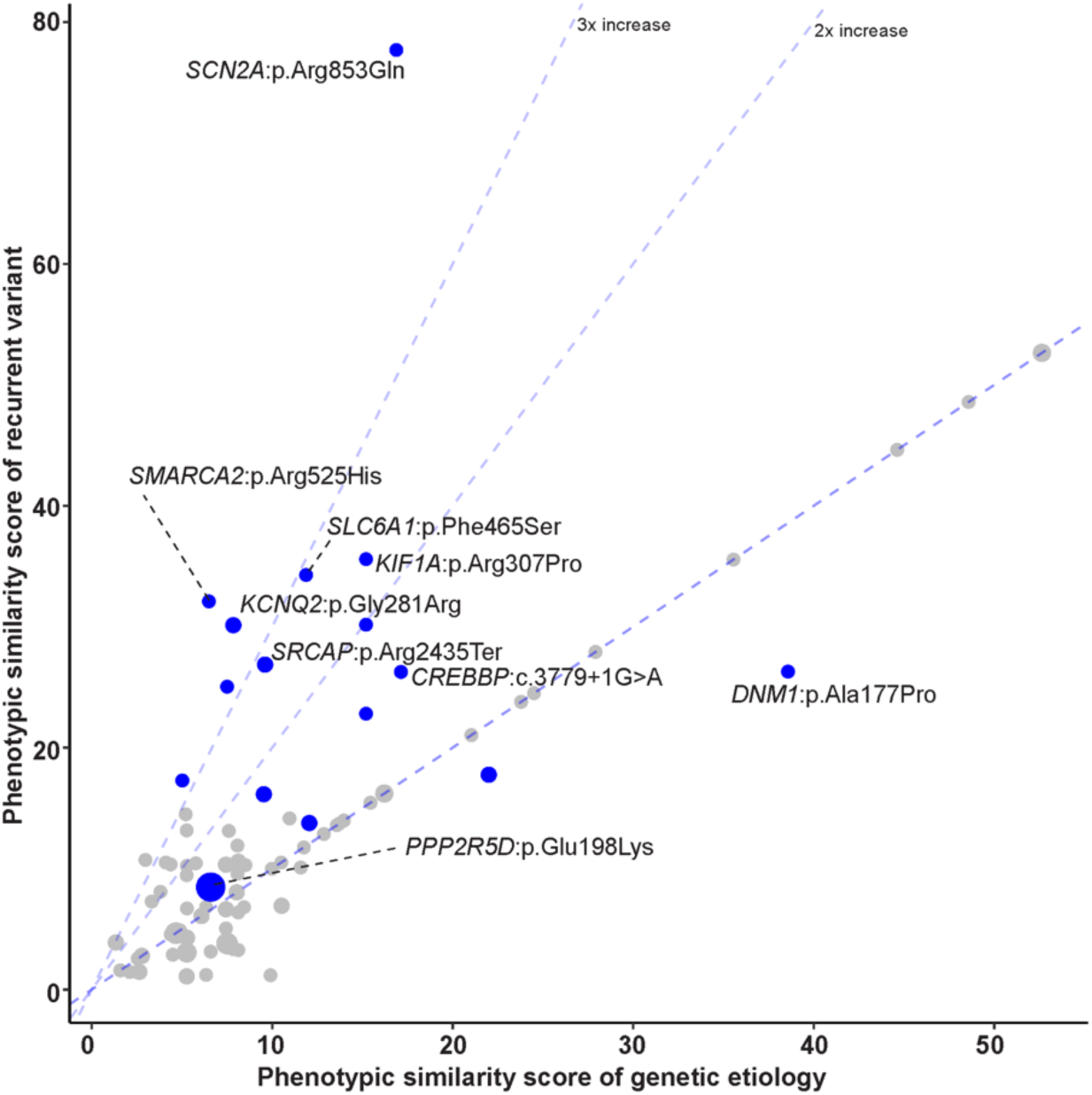
Phenotypic similarity of recurrent variants compared to genetic etiologies. Recurrent variants associated with specific genetic etiologies indicated higher median phenotypic similarity scores compared to the overall median similarity score of the respective genetic etiology. The dot size represents the number of individuals with the variant, blue dots indicate variants that reached statistical significance based on phenotypic evidence.

Recurrent variants with increased phenotypic similarity compared to gene-level similarity can be conceptualized as “diseases within diseases”, each with distinct features hinting at distinct molecular mechanisms. For example, *SCN2A*:p.Arg853Gln is known to demonstrate unique electrophysiological properties affecting the Nav1.2 sodium channel^30^, highlighted by HPO terms such as EEG with focal sharp waves (HP:0011196) and Infantile spasms (HP:0012469).

### Patterns of phenotypic similarities hint at hidden subgroups within genes

To examine whether genes for neurodevelopmental disorders encompass distinct subgroups based on phenotypic characteristics, we analyzed the distribution of pairwise similarity scores of each genetic etiology. We found that similarity scores varied widely across genes. Visual inspection of these distributions suggested possible subgroups within several genes (**Fig. 4a**).

**Fig. 4:**
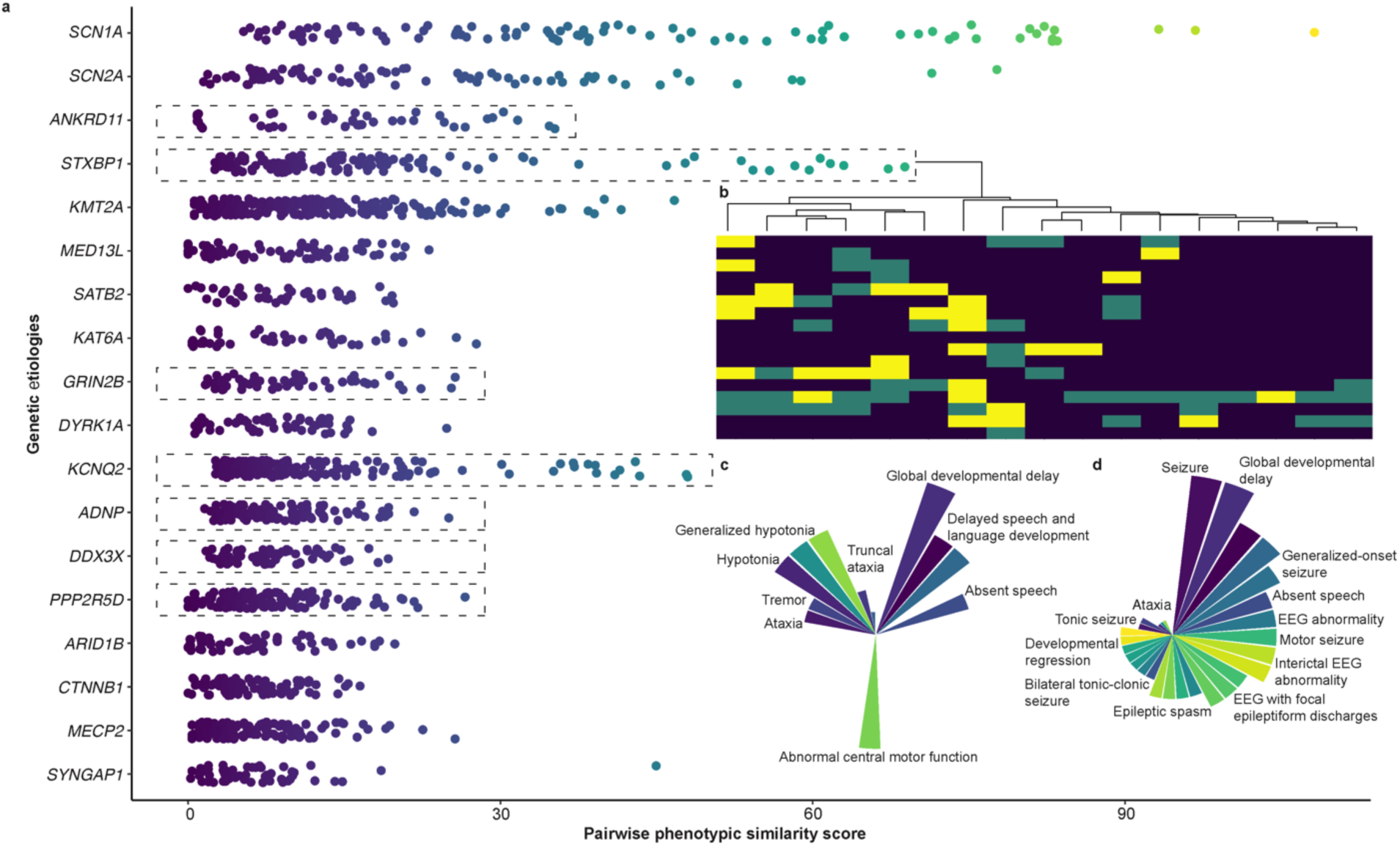
Phenotypic similarity patterns reveal subgroups within genetic etiologies. **a**, Pairwise phenotypic similarity scores for each genetic etiology suggest the presence of distinct subgroups within several genetic etiologies (indicated by the box). **b**, Focusing on the *STXBP1* gene, which exhibited a wide range of pairwise similarity scores, we performed clustering analysis that revealed two distinct subgroups. The heatmap represents hierarchical clustering, with the colors indicating phenotypic similarity patterns based on a weighted similarity score. The purple tile indicates higher phenotypic similarity, while yellow tiles indicate a lower similarity score. **c**, A radar plot highlighting hypotonia, ataxia, and tremor as more frequent in one group. **d**, Seizures, spasms, and abnormal EEG were more frequent in the other group.

Cluster analysis to formally define phenotypic subgroups revealed seven genes with distinct phenotypic clusters, including *STXBP1*, *ADNP*, *GRIN2B*, *DDX3X*, *PPP2R5D*, *ANKRD11*, and *KCNQ2*. For example, among individuals with *STXBP1*-related disorders (n=17), we identified one subgroup of individuals (n=7) all with seizures (HP:0001250), epileptic spasms (HP:0011097), and abnormal EEG findings (HP:0025373), while the second group (n=10) had muscular hypotonia (HP:0001252) and global developmental delay (HP:0001263) without seizures (**Fig. 4b-d**).

Knowledge of phenotypic subgroups is critical for the development of clinical outcome measures in a trial setting. Our results suggests that a large-scale phenotype analysis can aid in identifying hidden phenotypic subgroups, characterized either by increased phenotypic resemblance in individuals with recurrent variants or phenotypic clusters based on the distribution of pairwise comparisons.

### A phenotypic neighborhood approach identifies recognizable genes and clinical mimics

It is well established that individuals with a common genetic etiology may have highly similar features. Conversely, some groups of individuals display highly similar clinical features but have distinct or even unknown disease etiologies. We hypothesized that these “phenocopies”, or individuals with similar phenotypes lacking a common genetic etiology, could be detected by quantitatively identifying individuals who are clinically related to those with a particular genetic etiology.

We first analyzed the phenotypic neighborhood associated with individual genes. A phenotypic neighborhood consisted of individuals who shared phenotypic features and had high pairwise similarity scores. These individuals are connected by edges when a given individual is among the *k* most phenotypically similar to another. Initially, we assessed the 10-neighborhoods (i.e. *k*=10) of *SCN1A*, the gene which exhibited the highest overall phenotypic similarity in our dataset. This neighborhood included 61 individuals, including the 14 with known *de novo* variants in *SCN1A*. There were 29 individuals who had no identifiable *de novo* variants, among the 18 other individuals the most common non-*SCN1A* genes among this neighborhood were *DNM1* (n=2), and *GABRB3* (n=2). The identification of *de novo* variants in these genes in individuals who have highly similar clinical presentations to individuals with *SCN1A* variants suggests that some individuals with these genetic diagnoses may represent potential phenocopies of *SCN1A*-related disorders in the clinical setting, also highlighting the importance of a broader approach to genetic diagnosis.

Next, we explored which shared clinical features were the primary drivers of clinical relatedness within *SCN1A* and its broader phenotypic neighborhood (k=1000; **Fig. 5**). We verified that individual phenotypes associated with *SCN1A* remained statistically significant when compared to the larger neighborhood of individuals, often demonstrating even stronger associations within the 10-neighborhood and 100-neighborhood. Clinical features such as bilateral tonic-clonic seizures (HP:0002069; 61%), generalized myoclonic seizures (HP:0002123; 52%), and generalized clonic seizures (HP:0010818; 46%) were prevalent in this neighborhood (**Fig. 6a**), emphasizing that the other genes in the neighborhood of *SCN1A* causing generalized epilepsy with multiple seizure types can be phenotypic mimickers of *SCN1A*-related disorders, an observation that is often noted in the clinical care of individuals with genetic generalized epilepsies^31^. Moreover, this pattern of stronger associations suggests that including individuals with high phenotypic similarities to those with *SCN1A* increases the potential to identify key phenotypes of *SCN1A*-related disorders. This finding highlights the role of phenotypic neighborhoods in amplifying the essential features of a genetic disorder.

**Fig. 5:**
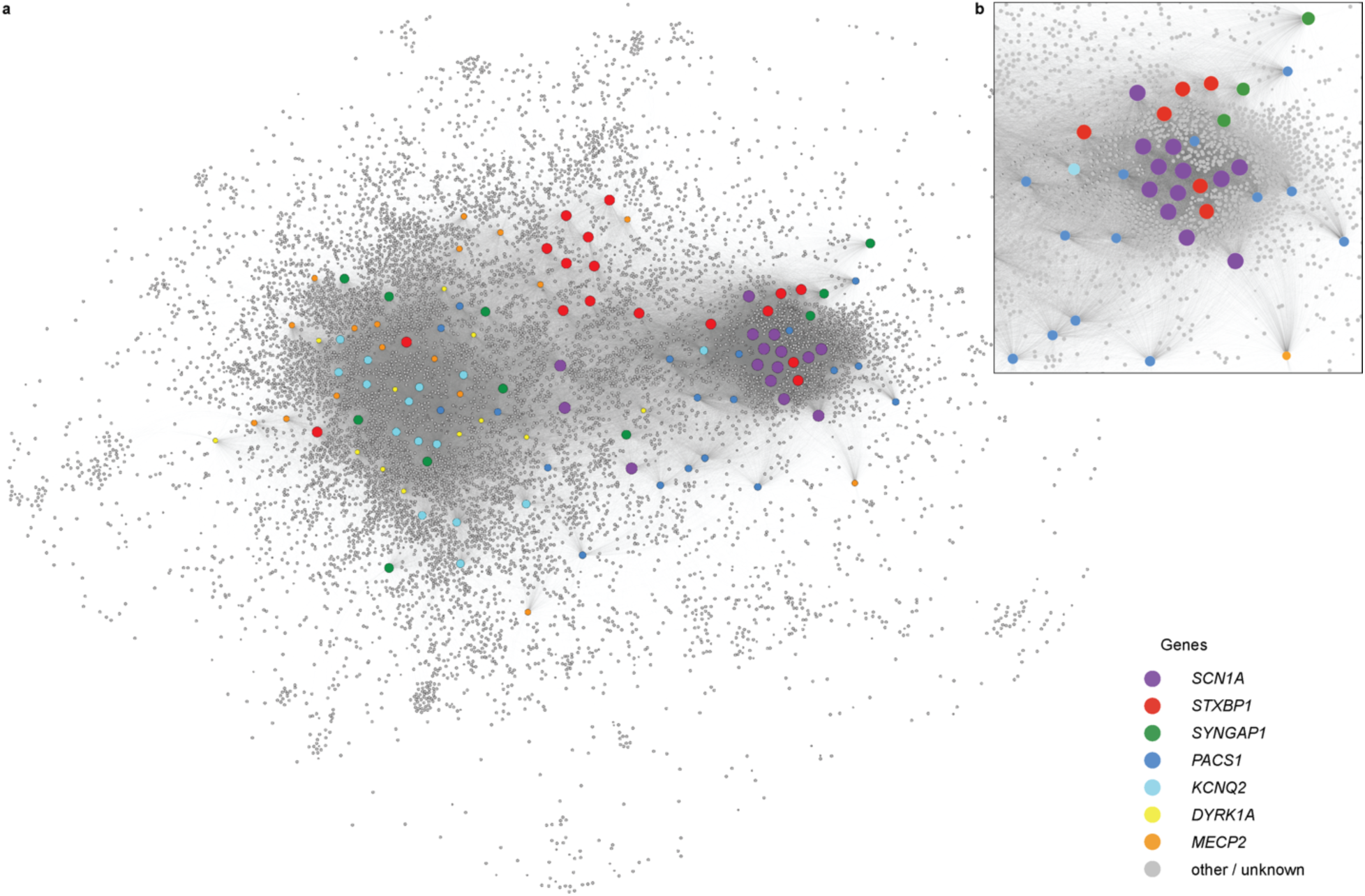
Network representation of phenotypic similarity in genetic neurodevelopmental disorders. **a**, A network visualization of individuals with *de novo* variants in known epilepsy genes and the top 1000 most phenotypically similar individuals. The distance between each individual is based on the phenotypic similarity score, computed using Fruchterman-Reingold algorithm in Gephi. Each node represents an individual, with colors and size of the dots indicating specific genetic etiologies. **b**, Focusing on the denser cluster reveals that individuals with *de novo* variants in *SCN1A* gene exhibit high phenotypic similarity to each other, suggesting a strong shared clinical presentation.

**Fig. 6:**
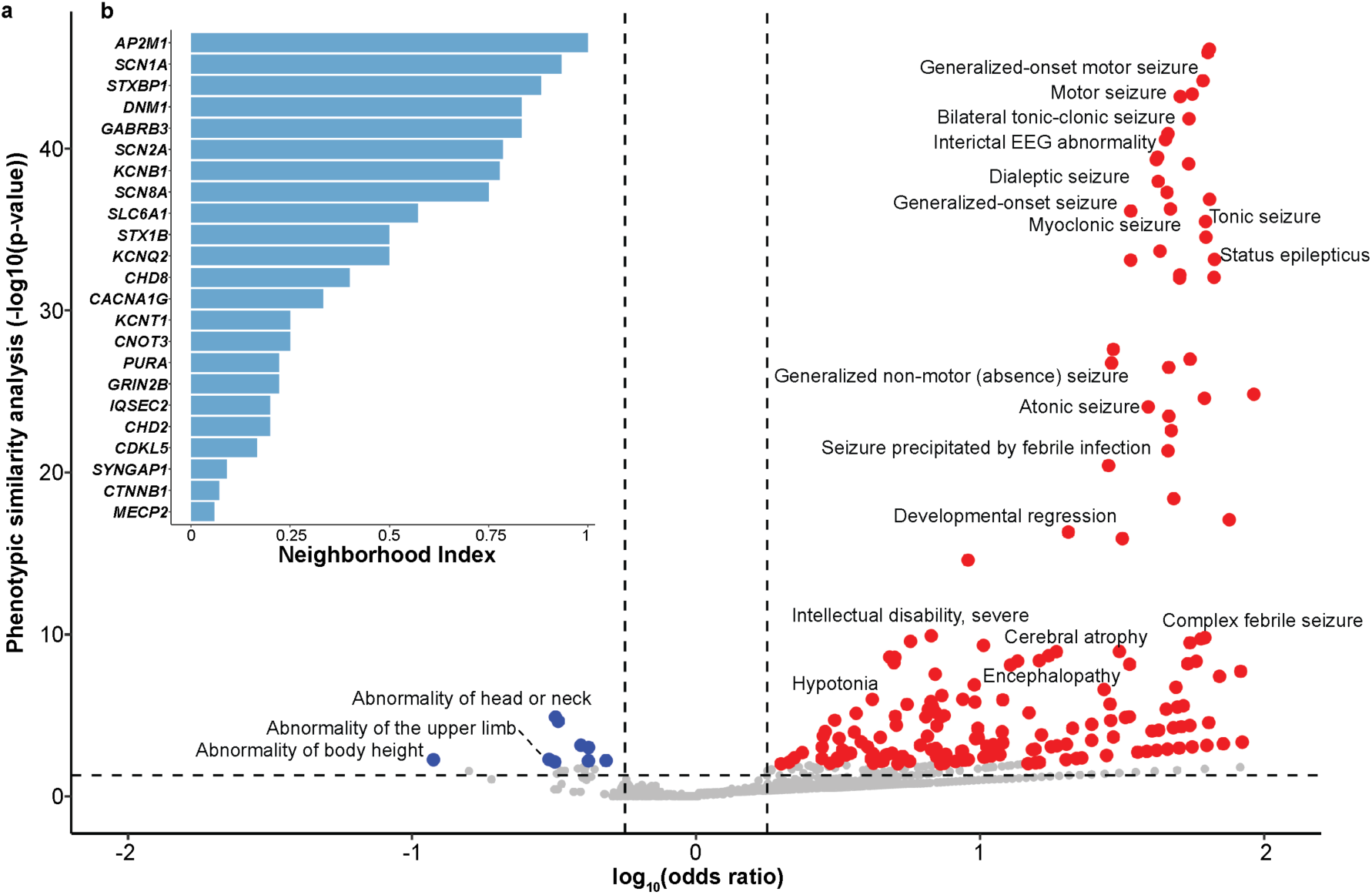
Phenotypic association in the *SCN1A* neighborhood. **a**, Volcano plot highlighting the HPO terms in individuals within the *SCN1A* neighborhood (k=100). The x-axis represents the odds ratio in log scale, indicating the strength of the association, while y-axis represents the statistical significance. Red dots indicate phenotypic terms positively associated with *SCN1A* such as Bilateral tonic-clonic seizure, Status epilepticus, and Febrile seizures, while blue dots indicate negative association with the phenotypes. **b**, Distribution of twin index for individuals within k=100 neighborhood, across all genetic etiologies.

We further hypothesized that computational detection of phenocopies through phenotypic neighborhoods could provide insight into the clinical recognizability of a genetic etiology. This was accomplished by capturing the estimated likelihood that two clinically similar individuals share the same genetic etiology, a measure we refer to as the “neighborhood index”. The neighborhood index represents the proportion of individuals within a subgroup who have at least one “genetic twin”, i.e., another individual with the same genetic etiology within the *k*-neighborhood. As expected, the neighborhood index increases with increasing values of *k*, reflecting the fact that including more individuals increases the likelihood of identifying someone with a *de novo* variant in the same gene. However, genes displayed distinct patterns of neighborhood indices. Notably, we found that *SCN1A* had a stable, high neighborhood index ranging from 78% to 100% across different *k* values (k=10,100,1000), whereas *SYNGAP1* had a low neighborhood index of 10% at *k*=50, which increased continuosly to 72% at *k*=1000 (**Fig. 6b**).

These results suggest that individuals with clinical presentations similar to those with *SCN1A*-related disorders are highly likely to possess a *de novo* variant in *SCN1A*, whereas the confidence for identifying an individual clinically similar to one with a *de novo* variant in *SYNGAP1* is lower. These findings are consistent with the established, consistent clinical presentation of individuals with *SCN1A*-related disorders, in contrast to the phenotypic heterogeneity among those with *SYNGAP1*-related disorders.

## Discussion

Both the genetic and phenotypic landscapes of neurodevelopmental disorders are highly complex, comprising hundreds of genetic etiologies and thousands of potential clinical features. Large-scale genomic sequencing has been the key technology in the discovery of genes associated with neurodevelopmental disorders. However, analysis of phenotypic data at a similar magnitude has not yet been performed. With precision therapies under development for a wide range of genetic conditions, understanding the link between genetic causes and linked clinical features is increasingly relevant.

Assessing clinical data at the same level as genomic information derived from massive parallel sequencing studies is challenging. In our dataset of 11,125 individuals with genomic and phenotypic data, we used the Human Phenotype Ontology (HPO) to systematically map clinical information onto a joint framework. The HPO represents a useful scaffold for this analysis for several reasons. First, HPO-based coding is already used in a wide range of studies and datasets. For example, the DDD cohort, which represents the largest cohort within our study, has been annotated phenotypically using HPO terms at the point of data collection^18^. Second, the HPO represents a simple yet comprehensive dictionary with a defined relationship between categories. Even though the HPO comprises >13,000 phenotypic categories, the relationship between phenotypes is limited to hierarchical relationships, enabling straightforward inference of broader phenotypic categories. In our cohort, phenotypic inference or “propagation” increased the median number of clinical terms per individual from 6 to 41 terms, a more than six-fold increase of clinical information. Finally, methods to assess phenotypic similarity have been extensively tested using HPO-coded phenotypes^6,10,12,25^.

The phenotypic depth in our dataset comprised 5,707 categories, with almost half of all clinical categories only present in two individuals or less. This observation reflects a common feature of clinical information in neurodevelopmental disorders: phenotypic data is sparse and highly dimensional^10^. However, we find that the size of the dataset compensates for its sparsity. For example, we were able to validate known genotype-phenotype associations, such the association of *de novo* variants in *KMT2A* with short stature (HP: 0004322; pval = 6.07 x 10^-4^), growth delay (HP:0001510; pval = 3.12 x 10^-4^), and hypertrichosis (HP:0000998; pval = 1.19 x 10^-6^), reflecting the well-established clinical spectrum of Wiedemann-Steiner syndrome^32^. Similarly, in individuals with *de novo* variants in *KCNQ2*, we found strong associations with generalized-onset seizures (HP:0002197; pval= 5.72 x 10^-6^), hypotonia (HP:0001252; pval= 1.52x 10^-2^), and EEG with burst suppression (HP:0010851; pval = 2.15 x 10^-7^), known clinical features linked to *KCNQ2*-related disorders^33^.

In order to assess the phenotypic gestalt in neurodevelopmental disorders more holistically, we focused on phenotypic similarity, a measure that determines clinical relatedness based on all phenotypic features of an individual. We identified 69 phenotypically significant genes, including established neurodevelopmental genes including *SCN1A*, *SCN2A*, *SCN8A*, *AP2M1*, *DNM1*, *GABRB3*, and *STX1B*^19,34^. We did not observe higher than expected phenotypic similarity for synonymous *de novo* variants, emphasizing that phenotypic similarity algorithms are robust and do not artificially inflate statistical significance. This finding suggests that a large number of neurodevelopmental genes have strong phenotypic fingerprints, which may be critical to identify disease-specific outcome measures in clinical trials.

We found that 15 phenotypically similar genes did not achieve genomic significance. These genes represent 5.7% of the 261 genes with *de novo* variants in two or more individuals and account for 33 individuals, 3.3% of the 992 individuals impacted by these 261 genetic etiologies. This increase in individuals with explained genetic causes emphasizes the utility of phenotypic similarity algorithms as a complementary means to identify neurodevelopmental genes. These methods may be particularly relevant given that several hundred neurodevelopmental genes are yet to be identified^23^, and approaches based on genomic information alone are expected to have diminishing returns in large and heterogeneous datasets^22^.

We observed two unexpected findings when we examined subgroups defined by genetic or phenotypic features. First, we found that the phenotypic landscape of recurrent *de novo* variants is inherently complex. For almost half of all recurrent *de novo* variants (35/81, 43%), phenotypic similarity exceeds gene-level similarity. Moreover, recurrent variants *SMARCA2*, *PPP2R5D*, and *TCF7L2* are phenotypically similar despite lack of clinical resemblance when assessing all *de novo* variants in these genes. This finding suggests that recurrent *de novo* variants may have unique biological consequences that frequently differ from the disease mechanism of the wider genetic etiology. Second, when analyzing phenotypic subgroups based on pair-wise clinical resemblance, we observed phenotypic clusters in seven neurodevelopmental genes, including several genes previously thought to result in homogeneous phenotypes. Defining meaningful subgroups is a key prerequisite for future clinical trials, and our results suggest that large clinical and genomic datasets may provide an initial insight for the potential existence of novel subgroups in well-known neurodevelopmental genes.

Many individuals with genetic neurodevelopmental disorders present similarly to individuals with different or undetectable genetic diagnoses. To explore how neurodevelopmental genes cluster within groups of individuals with related phenotypes, we introduced the concept of the neighborhood index as a novel concept. This index describes the likelihood that clinically similar individuals share the same neurodevelopmental gene. Accordingly, a high neighborhood index suggests a strong ability to recognize a genetic neurodevelopmental disorder in a clinical setting. We find that neighborhood indices vary widely across genes with strong phenotypic similarity, suggesting that only a small subset of neurodevelopmental genes including *AP2M1*, *SCN1A*, *STXBP1*, *DNM1*, and *GABRB3* share clinical features that allow these conditions to stand out from the wide phenotypic range seen in individuals with neurodevelopmental disorders.

In summary, by jointly assessing genomic and clinical features in neurodevelopmental disorders in 11,125 individuals with genomic and phenotypic data, we discover an unexpectedly complex disease landscape. As genomic and phenotypic data continues to grow, exploring the interplay of clinical and genomics features may provide novel insights into phenotypic fingerprints linked to neurodevelopmental genes and patterns within subgroups defined by genetic or clinical features.

## Methods

### Genetic analysis

All exome sequencing data from EPGP, EuriEPINOMICS-RES, and our local CHOP trio cohort were returned in FASTQ format. All the data were re-analyzed using a standardized pipeline. Raw sequencing reads were mapped to the human reference genome HS37d5 using Burrows-Wheeler Alignment (BWA) MEM algorithm to produce an aligned BAM file^35^. Duplicate reads were removed using Picard, and base quality scores were recalibrated using GenomeAnalysisToolKit (GATK) software^36^ to account for the systematic bias by the sequencer. Variant calling (SNP and Indel) was performed using HaplotypeCaller and genotyped using GenotypeGVCFs in GATK. For the DDD dataset, we got trio-variant call files also aligned to the HS37d5 human reference genome. To make the dataset harmonized, all the variants from these cohorts were liftovered to human genome build 38.

The whole genome sequencing dataset we got access through the Birth Defects Biorepository, data processing were performed by the genomics platform at the Broad Institute of MIT and Harvard. DNA libraries were prepared using the Illumina Nextera or Twist exome capture (∼38 Mb target) and sequenced with 150 bp paired-end reads, achieving >85% of targets covered at 20x and a mean target coverage of >55x. Sequencing data were processed through a pipeline based on Picard, with read mapping performed using the BWA aligner to the human genome build 38 (GRCh38). Variants were called using the Genome Analysis Toolkit (GATK) HaplotypeCaller package version 3.5, following best practices for variant detection.

Variant annotation for all the datasets were done using ANNOVAR and VEP for expected functional sequence^21,37^. Other annotations also included allele frequencies from gnomAD and CADD (Combined Annotation Dependent Depletion) scores^38,39^. Individual genes were further annotated with Residual Variant Intolerance Score (RVIS) percentile, and pLI or probability of loss-of-function intolerance^40^. Variants were categorized as *de novo*, homozygous, or compound heterozygous if the following filtering criteria was fulfilled: genotyping quality in all samples were greater than 20, total read depth greater was than 10, the variant was not present in the gnomAD database, RVIS percentile was less than 55, and the gene displayed dominant inheritance in Online Mendelian Inheritance in Man (OMIM)^40^. Additionally, we computed the p-value for all genes with *de novo* variants using denovolyzeR which compares the expected number of *de novo* variants against the observed number of *de novo* variants per gene^24^.

### Similarity analysis

The Human Phenotype Ontology (HPO) provides a standardized biomedical representation of >15,000 phenotypes in human disease^15^. Every term in HPO represents a distinct clinical feature. All the individuals in our cohort were annotated with HPO terms capturing their clinical presentation. For the final analysis, all HPO terms were propagated using automated harmonization, as previously described, after removing obsolete and duplicate terms based on HPO release v2023-08-02. The frequency (f) of each HPO term was computed, allowing us to calculate Information Content (IC), defined as -log_2_(f).

We calculated phenotypic similarity between individuals using three different algorithms. First, Resnik similarity uses information content (IC) of a common parent term, also known as most informative common ancestor (MICA), for two individual HPO terms^41^. Pairwise similarity for two individuals is computed using the following, where *P_1_* and *P_2_* are two individuals in the cohort having *m* and *n* annotations respectively, and *S_ij_* is the information content of the MICA of HPO terms *i* and *j*:

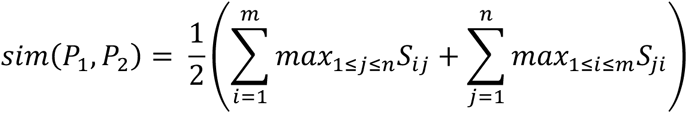

In the next method, we calculated the similarity of two individuals by sum of the IC of the shared or intersection of HPO terms after propagation. We refer to this method as the ‘cube’ algorithm, also known as total overlap:

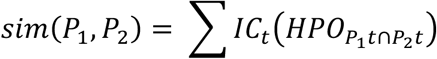

The last similarity method entailed computing the intersection or total overlap of two individuals after filtering out redundant or implied terms from each individual using the function minimal_set from package ontologyIndex in R:

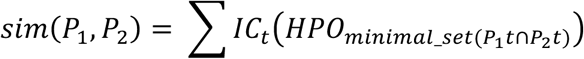

For the primary analysis we used the Resnik-mod (Equation 1) algorithm as it is the most well-established method. To test the significance of the phenotypic similarity within a particular gene with *n* individuals, we computed the distribution of median similarities of *n* individuals chosen randomly across one million permutations. We then compared this distribution to the median similarity of the respective gene with *n* individuals. Comparing the expected and observed median similarity resulted in an exact p-value.

### Clustering analysis

We used agglomerative clustering with Ward’s method within the cluster package in R to cluster individuals and clinical features within individual gene subgroups. For HPO features, we weighted terms by their information content prior to clustering. For individuals, we used the inverse of their Resnik phenotypic similarity as a measure of distance. We determined the number of meaningful clusters by minimizing the ratio of mean height within the cluster to height between clusters for all values between two and ten.

### Neighborhood analysis

We defined an individual’s phenotypic *k*-neighborhood as the individual along with the *k* most phenotypically similar individuals in our cohort. The *k*-neighborhood of an entire genetic etiology was then defined as the union of *k-*neighborhoods of all individuals within the gene subgroup. For example, the 10-neighborhood of a gene consisted of all individuals with a *de novo* variant in that gene, along with the 10 most similar individuals to each within the larger cohort.

Next, for each genetic cause, we measured the “neighborhood index.” We computed the *k*-neighborhood for each individual within a given gene genetic etiology and determined how many individuals had at least one *k*-neighbor with the same genetic cause. We computed neighborhood indices for each genetic etiology, for *k* values in increments of 10 up to 100, and k values of 500, and 1000 and compared twin indices across all genetic etiologies in our cohort.

**Extended Data Table 1.** Genes with most identified *de novo* variants across cohorts.

| **Gene** | **Ensembl gene ID** | **n** | **Recurrent variant** | **OMIM #** |
| --- | --- | --- | --- | --- |
| **KMT2A** | ENSG00000118058 | 22 | ENSP00000436786.2:p.Asn3383SerfsTer2 (n=2) | 159555 |
| **KCNQ2** | ENSG00000075043 | 20 | ENSP00000352035.2:p.Gly281Arg (n=3) | 602235 |
| **STXBP1** | ENSG00000136854 | 17 | ENSP00000362396.2:p.Arg367Ter (n=2) | 602926 |
| **MECP2** | ENSG00000169057 | 16 | ENSP00000395535.2:p.Arg145Cys (n=5) | 300005 |
| **PPP2R5D** | ENSG00000112640 | 16 | ENSP00000417963.1:p.Glu198Lys (n=12) | 601646 |
| **ADNP** | ENSG00000101126 | 15 | ENSP00000483881.1:p.Asn832LysfsTer81 (n=2) | 611386 |
| **CTNNB1** | ENSG00000168036 | 14 | ENSP00000344456.5:p.Arg535Ter (n=3) | 116806 |
| **SCN1A** | ENSG00000144285 | 14 | - | 182389 |
| **SCN2A** | ENSG00000136531 | 13 | ENSP00000364586.2:p.Arg853Gln (n=2) | 182390 |
| **DDX3X** | ENSG00000215301 | 13 | ENSP00000494040.1:p.Arg292Ter (n=3) | 300160 |
| **MED13L** | ENSG00000123066 | 12 | - | 608771 |
| **ARID1B** | ENSG00000049618 | 11 | ENSP00000490491.2:p.Glu1879AlafsTer9 (n=2) | 614556 |
| **DYRK1A** | ENSG00000157540 | 11 | ENSP00000494572.1:p.Arg246Ter (n=2) | 600855 |
| **GRIN2B** | ENSG00000273079 | 11 | ENSP00000477455.1:p.Gly689Ser (n=3) | 138252 |
| **SYNGAP1** | ENSG00000197283 | 11 | - | 603384 |
| **ANKRD11** | ENSG00000167522 | 10 | - | 611192 |
| **KAT6A** | ENSG00000083168 | 10 | - | 601408 |
| **SATB2** | ENSG00000119042 | 10 | ENSP00000401112.1:p.Arg399His (n=2) | 608148 |
| **ASXL3** | ENSG00000141431 | 9 | ENSP00000269197.4:p.Arg1444Ter (n=3) | 615115 |
| **KCNB1** | ENSG00000158445 | 9 | - | 600397 |
| **CDKL5** | ENSG00000008086 | 8 | - | 300203 |
| **GATAD2B** | ENSG00000143614 | 8 | ENSP00000357644.4:p.Arg116Ter (n=3) | 614998 |
| **KAT6B** | ENSG00000156650 | 8 | - | 605880 |
| **IQSEC2** | ENSG00000124313 | 7 | - | 300522 |
| **KIF1A** | ENSG00000130294 | 7 | ENSP00000438388.1:p.Arg307Pro (n=2) | 601255 |
| **NEXMIF** | ENSG00000050030 | 7 | - | 300524 |
| **SCN8A** | ENSG00000196876 | 7 | - | 600702 |
| **SETD5** | ENSG00000168137 | 7 | ENSP00000385852.2:p.Arg768Ter (n=3) | 615743 |

**Extended Data Table 2.** Comparison of phenotypic similarity among recurrent *de novo* variants and associated genetic etiologies.

| Gene | No. of individuals | median similarity of Gene | Recurrent variant | No. of individuals recurrent | median similarity of variant | distinct phenotypic features |
| --- | --- | --- | --- | --- | --- | --- |
| **PPP2R5D** | 16 | 6.6 | p.Glu198Lys | 12 | 8.5 | Generalized Hypotonia, Macrocephaly |
| **KCNQ2** | 20 | 7.8 | p.Gly281Arg | 3 | 30.1 | Encephalopathy, Generalized tonic seizure, EEG with generalized epileptiform discharges |
| **GATAD2B** | 8 | 9.5 | p.Arg116Ter | 3 | 16.1 | Neonatal hypotonia, Hypoplasia of the corpus callosum |
| **SRCAP** | 6 | 9.6 | p.Arg2435Ter | 3 | 26.9 | Growth abnormality, short stature, Frontal bossing |
| **KCNQ3** | 5 | 12.1 | p.Arg230Cys | 3 | 13.8 | Atonic seizure, EEG with focal sharp waves, Facial asymmetry |
| **ADNP** | 15 | 7.5 | p.Asn832LysfsTer81 | 2 | 25.1 | Microcephaly, axial hypotonia, coloboma |
| **SCN2A** | 13 | 16.9 | p.Arg853Gln | 2 | 77.7 | Febrile seizure (within age range of 3 months to 6 years), atypical absence seizure, EEG with spike-wave complexes (<2.5 Hz) |
| **DNM1** | 6 | 38.6 | p.Ala177Pro | 2 | 26.3 | Generalized myoclonic seizure, Interictal epileptiform activity |
| **KIF1A** | 7 | 15.2 | p.Arg307Pro | 2 | 35.6 | Intellectual disability, spasticity, Hypertonia |
| **KIF1A** | 7 | 15.2 | p.Arg307Gln | 2 | 30.2 | Neonatal hypotonia, Severe global developmental delay |
| **KIF1A** | 7 | 15.2 | p.Pro305Leu | 2 | 22.8 | Seizure, moderate global developmental delay |
| **SLC6A1** | 6 | 11.9 | p.Phe465Ser | 2 | 34.3 | Growth delay, Bilateral tonic-clonic seizure with focal onset |
| **CREBBP** | 5 | 17.1 | c.3779+1G>A | 2 | 26.3 | Growth delay, abnormality of cardiovascular system |
| **TCF7L2** | 4 | 5.1 | c.686-1G>A | 2 | 17.3 | Febrile seizure (within age range of 3 months to 6 years), Talipes |
| **SMARCA2** | 4 | 6.5 | p.Arg525His | 2 | 32.1 | Wide mouth, prominent nasal bridge |

**Extended Data Fig. 1:**
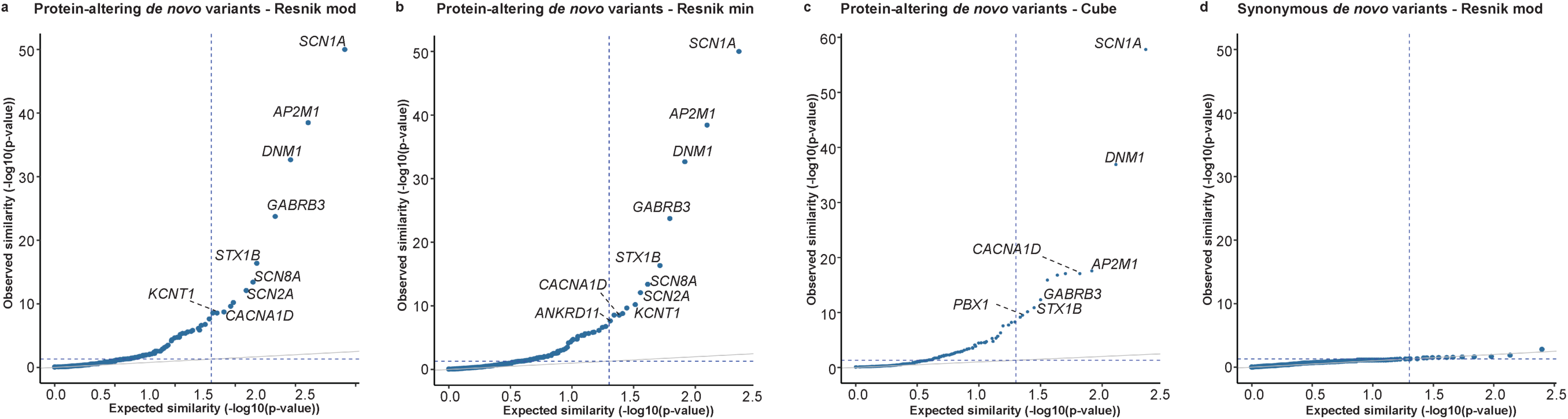
Phenotypic similarity across different algorithms. **a-c**, QQplots showing us the most significant gene based on phenotypic similarity analysis using HPO terms with three different algorithms. **d**, The observed phenotypic similarity of synonymous *de novo* variants remained low, validating the findings in our methods.

## Acknowledgements

The authors thank the participants and their family members for taking part in this study. We would like to thank or clinical research coordinators for their extensive support in enrolling research participants and admisitrative assistance. The DDD study presents independent research commissioned by the Health Innovation Challenge Fund [grant number HICF-1009-003]. This study makes use of DECIPHER (http://www.deciphergenomics.org), which is funded by Wellcome [grant number WT223718/Z/21/Z]. See Nature PMID: 25533962 or www.ddduk.org/access.html for full acknowledgement. The CHOP Birth Defects Biorepository is supported by the National Center for Advancing Translational Sciences, National Institutes of Health, through Grant UL1TR001878.

## Author contributions

I.H. and S.G. conceived the project. S.G., P.D.G, and S.M.R. were responsible for data curation. S.G, P.D.G, and S.P. developed the methods for the analysis. S.G, P.D.G, S.P., and S.R.M. conducted the formal analysis. S.G., S.P.,S.R.C., and I.H. drafted the manuscript. All authors reviewed and edited the manuscript.

## Ethics declaration

Informed conset for participation in this study was obtained from the parents of all probands in our CHOP EGRP cohort. The study was completed per protocol with local approval by CHOP Institutional Review Board (IRB 15-12226).

## Data and code availability

Datasets in a deidentified format will be available upon request to the corresponding author. The DDD dataset files are publicly available from the European Genome-phenome Archive (study ID: EGAS00001000775), and the EPGP dataset is available from dbGap (accession: phs000653.v1.p1) respectively. Code for the primary analysis is available at https://github.com/helbig-lab/trio_hpo

## Additional information

Correspondence and request for materials should be addressed to Ingo Helbig. Reprints and permissions information is available at http://www.nature.com/reprints.

